# MinYS: Mine Your Symbiont by targeted genome assembly in symbiotic communities

**DOI:** 10.1101/2019.12.13.875021

**Authors:** Cervin Guyomar, Wesley Delage, Fabrice Legeai, Christophe Mougel, Jean-Christophe Simon, Claire Lemaitre

**Affiliations:** Univ Rennes, CNRS, Inria, IRISA - UMR 6074, F-35000 Rennes, France; INRA, UMR 1349 INRA/Agrocampus Ouest/Université Rennes 1, Institut de Génétique, Environnement et Protection des Plantes (IGEPP), F-35653 Le Rheu, France; iDiv – German Centre for Integrative Biodiversity Research, Deutscher Platz 5e, D-04103 Leipzig, Germany

## Abstract

Most metazoans are associated with symbionts. Characterizing the effect of a particular symbiont often requires to get access to its genome, which is usually done by sequencing the whole community. We present MinYS, a targeted assembly approach to assemble one particular genome of interest from such metagenomic data. First, taking advantage of a reference genome, a subset of the reads is assembled into a set of backbone contigs. Then, this draft assembly is completed using the whole metagenomic readset in a *de novo* manner. The resulting assembly is output as a genome graph, allowing to distinguish different strains with potential structural variants coexisting in the sample. MinYS was applied to 50 pea aphid re-sequencing samples, with low and high diversity, in order to recover the genome sequence of its obligatory bacterial symbiont, *Buchnera aphidicola*. It was able to return high quality assemblies (one contig assembly in 90% of the samples), even when using increasingly distant reference genomes, and to retrieve large structural variations in the samples. Due to its targeted essence, it outperformed standard metagenomic assemblers in terms of both time and assembly quality.

## Introduction

The advances of molecular techniques revealed the importance of micro-organisms in every ecosystem. In particular, it is now well acknowledged that most metazoans are associated with microbes, forming complex entities named as holobionts. In these systems, microbes are interacting together as well as with their host, and host-microbe interactions can have significant effects on the host phenotype (1). Many animals have microbial associations with one ore a few specific symbionts. This is for example the case for corals associated with the algal symbiont *Symbiodinium* (2), squids with *Vibrio* (3), woodlice with *Wolbachia* (4) or hemipteran insects with specific obligatory and facultative bacterial symbionts (5). As there is usually no way to target the DNA of one or several particular symbionts for sequencing, the whole community is usually sequenced, resulting in a metagenomic dataset mixing host and symbiont reads. These datasets are unbalanced: the great majority of the reads often originate from the host genome, but since the genomes of the symbionts are often several orders of magnitude smaller than the one of the eukaryotic host, symbiont genomes can have large read depth in such samples. This allows to extract relevant information about the symbionts, but requires significant effort, since the host reads are a computational burden for most analyses. In that context, providing bioinformatic tools that allow to assemble a particular genome of interest from a metagenomic sample, ignoring the overwhelming amount of reads from other organisms, is a key question to better characterize symbiont genomes, and therefore decipher particular host-symbiont relationships.

In most cases, some knowledge and genomic resources about the symbiont of interest are already available. And a common problem is to recover the full genomic sequence of the particular symbiont present in the metagenomic dataset, using available genomic resources. The way we address this problem depends on the availability of a reference genome and its closeness to the considered species. In the easiest but rarest case, a reference genome of good quality is available for the considered species, and a mapping-based approach is classically performed. Reads from the whole metagenomic datasets are mapped on this reference genome and small variants are called to characterise the strain at play. This approach does not output *per se* a genome sequence but rather a list of punctual variants with respect to a reference one, without any evidence that this list is exhaustive and ignoring potential larger structural differences, such as large novel insertions or deletions. This approach may therefore miss crucial genomic information even where a very close reference genome is available, since it is well known that bacterial strains can have highly variable accessory genomes, eventually responsible for pathogenicity or other phenotypic effects (6).

To circumvent these drawbacks, a more classical approach relies on *de novo* genome assembly to obtain full genomic sequences. In this metagenomic context, the whole community is assembled and available genomic resources are used to select afterwards among the assembled contigs the ones originating from the species of interest. Assembling genomes from metagenomic data is a challenging task, because of both the many species coexisting in the samples and the polymorphism within these species. Although many recent software are devoted to this task (7–9), metagenomic assemblies are often very fragmented and come with a high computational cost (10). In this particular case, where reads originating from the targeted organism are a minority in the whole metagenomic read set, this computational cost seems unnecessary and by focusing on a subsample of reads we could hope to obtain less fragmented assemblies.

Therefore, combining mapping-based and assembly-based approaches in a so-called targeted genome assembly seems a promising way to facilitate the study of such communities. In such approaches, a subset of the metagenomic reads is first recruited by mapping to a reference genome and then de novo assembled. The relatedness of the used genome is a key parameter of the approach, and methods have to be able to incorporate non recruited reads in the assembly, in order not to miss divergent or strain-specific regions.

Several tools, such as MITObim (11), LOCAS (12), Pilon (13) or IMR/DENOM (14), were designed following this idea, combining reference alignment and *de novo* assembly. However, all of them show some limitations because of which they appear not adapted to metagenomic data. Primarily, they all work under the assumption of genomic homogeneity (no polymorphism), which is very rarely met in the metagenomic context. They can therefore not return coexisting strains. Moreover, the architecture of the reference genome is used as a starting-point for the assembly in most of the cases, and they are unable to deal with large structural variants with respect to the reference genome (IMR/DENOM, Pilon, LOCAS). Finally, some tools are designed either for short genomes or small resequencing datasets and they are not scaling up with the size of metagenomic datasets (MITObim, LOCAS).

To our knowledge, a single tool has been designed to guide the assembly by a reference genome in a metagenomic context. MetaCompass is a pipeline described in a pre-print (15), that 1) recruits a subset of reads by mapping on a reference genome, 2) assembles them into contigs and 3) assembles all the remaining non-recruited reads to recover the regions potentially missing in the reference genome. This last step amounts at the end to assemble the whole community instead of one single genome of interest, making this approach reference-guided rather than targeted.

In this work, we present MinYS, a novel method for the assembly of a genome of interest from metagenomic data, that does not require to assemble the full readset. This method can recover large regions absent from the given reference genome, it makes no assumption on the order or orientation of the regions homologous with the reference and it is able to return several different solutions reflecting the potential strain diversity inside the sample. It is based on two main steps, a reference based recruitment and assembly of reads, followed by a targeted assembly using the whole readset, filling the gaps between the contigs assembled beforehand.

We applied this method to reconstruct genomes from 50 metagenomic samples of the pea aphid *Acyrthosiphon pisum*. Focusing on the primary endosymbiont *Buchnera aphidicola*, we demonstrated the ability of MinYS to assemble complete bacterial genomes in a single contig using a remote reference genome as a primer, even when structural variability is present.

## Material and Methods

### A novel method for targeted genome assembly with metagenomic data

#### Approach overview

The method described in this work relies on a two-step pipeline, described in Figure 1.

**Fig. 1.**
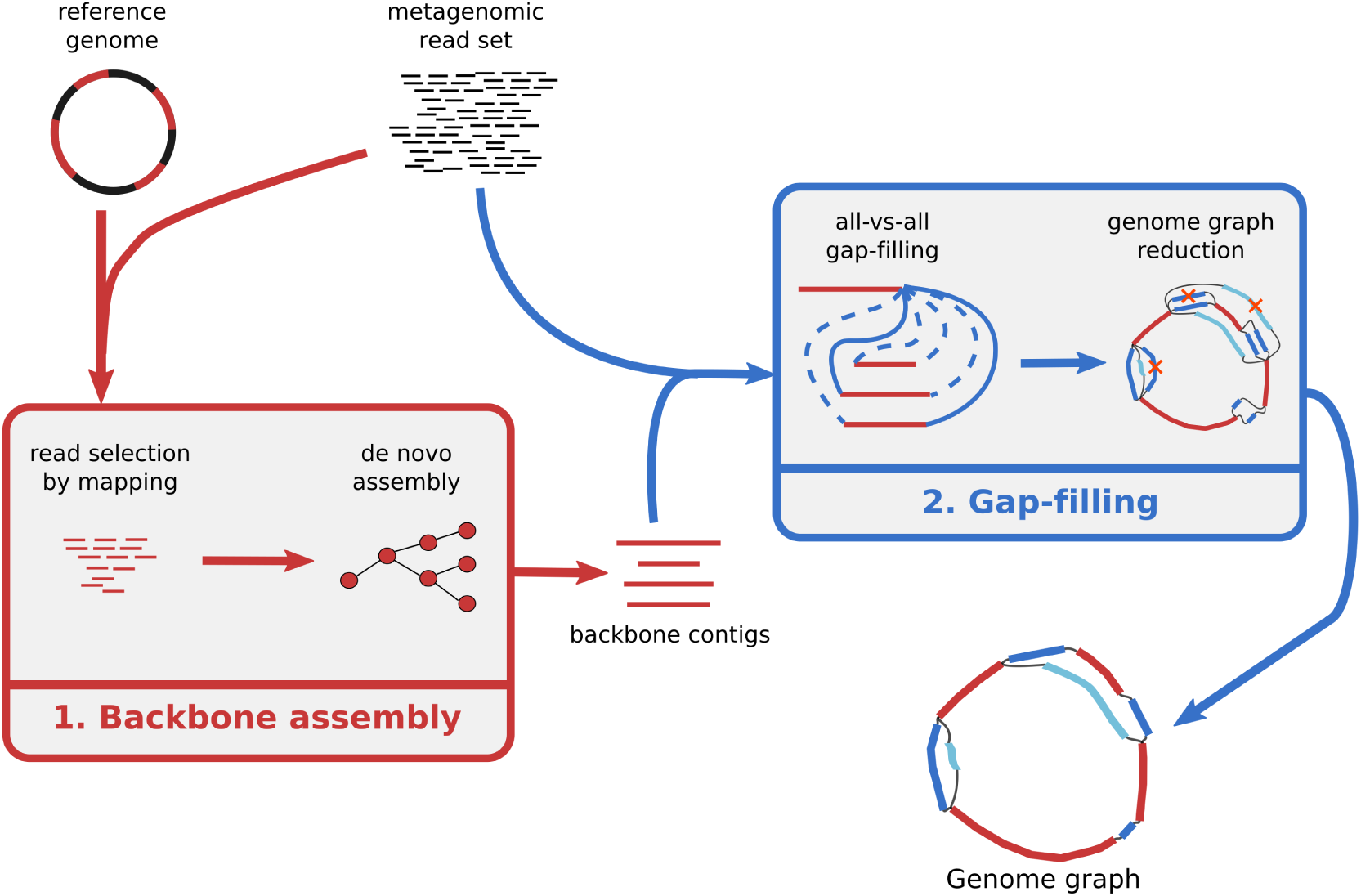
Overview of MinYS approach, for targeted genome assembly. MinYS takes as input a reference genome and a set of metagenomic reads, and outputs the targeted genome assembly as a genome graph encompassing the potential strain diversity contained in the sample.

The first step uses a given reference genome to build an incomplete but trustworthy assembly from a subset of the input metagenomic reads, here-after called the backbone contigs. The second step uses the whole set of metagenomic reads to extend the previously assembled contigs and form a complete assembly, without any *a priori* on the order and orientation of the backbone contigs. The result of the pipeline is a genome graph encompassing the stuctural diversity detected on the assembled genome. This graph can be exploited by extracting contigs, or paths of the graph that represent different strains.

#### Assembly of backbone contigs

The first step requires a metagenomic readset and a reference genome, and returns contigs that are *de novo* assembled using only reads mapped on the reference genome. All metagenomic reads are mapped against the reference genome using BWA MEM (16), and the mapped reads are kept and *de novo* assembled using the Minia (17) assembler. Although any assembler can be used in this step, we use Minia for its low memory footprint, and its assembly algorithm similar to the one used in the second step of the method. The amount of mapped reads and the size and contiguity of the resulting assembly depend on the sequence similarity between the reference genome and the targeted genome. However, the goal of this step is not to produce a complete assembly of the target genome but rather to generate high quality contigs, that can reliably be used for the upcoming gap-filling step. To ensure this, we set up Minia with more stringent parameters than for a usual assembly task (ie. with a higher kmer size and a higher minimal abundance threshold). In the same goal, only contigs larger than a user-defined threshold (400 bp by default) are kept for the second step.

#### Gap-filling between backbone contigs

The essential step of the pipeline is the gap-filling between backbone contigs, which enables the assembly of regions of the targeted genome that are absent or too much divergent in the reference genome. This is made possible by a targeted assembly of the whole readset using the backbone contigs as primers. This step does not require any relative ordering or orientation of contigs, since all possible combinations are tested during gap-filling. As a result, structural variants can be detected, either compared to the reference genome or within the sample.

This step is based on a module of the software *MindTheGap*, originally developed for the detection and assembly of long insertion variants (18). MindTheGap is built upon the GATB library (19), that offers memory and time efficient data structures for De Bruijn graphs. The *fill* module of *MindTheGap* builds a de Bruijn graph of the entire input read set, and performs a local assembly between the left and the right kmers of each insertion site, by looking for all the paths in the De Bruijn graph starting from the left (source) kmer and ending in the right (target) kmer. In this work, we took advantage of this module of *MindTheGap* and adapted it to the problem of simultaneous gap-filling between multiple contigs. It has been modified to make possible the gap-filling between a source kmer and multiple target kmers, enabling the “all versus all” gap-filling within a set of contigs with a linear time increase. The improvements that have been made to MindTheGap in the context of MinYS are available as a new option of MindTheGap (named *contig mode*) and are presented in Supplementary Section 1.

MindTheGap gap-filling results are output as a sequence graph in the GFA format (Graphical Fragment Assembly), containing all input contigs and their successful gap-filling sequences as nodes, together with their overlap relationships as edges.

#### Graph simplification

By construction, the raw sequence graph output by MindTheGap is likely to contain redundant sequence information. A graph simplification step has been therefore developed to remove uninformative sequence redundancy and hence ease the analysis and usage of such a genome graph. The different steps of this process are represented in Fig S2 of the supplementary material.

First, it is likely that two contigs are linked in the graph by two gap-fillings with reverse-complement sequences, one starting from the left contig and the other one starting from the right contig in the reverse orientation. Such reciprocal links are removed, when their sequence identity is over a 95% identity threshold.

Secondly, when several gap-filling sequences start (or end) from the same seed contig, it is possible that a subset of them have an identical prefix (resp. suffix) and start to diverge only after a potential large distance from the seed (resp. target) contig. This results in redundant sequence information in the final sequence set. A node merging algorithm is applied, in order to return a final set of sequences (nodes of the graph) that do not share large identical subsequences as prefix or suffix. Sets of sequences sharing the same prefix of size *l* are built (*l* equals 100 by default). Within each set, the sequences are then compared to find the first divergence between all sequences. A new node is added to the graph, containing the repeated portion of the sequences, and repeated nodes are shortened accordingly. This process is applied iteratively to every node, including the newly created nodes, for which a subset of neighbors may still show identical sequences.

Finally, simple linear paths, with no branching nodes, are merged into a single node.

After the simplification process, the graph may not be a linear sequence because of intra-sample polymorphism or assembly uncertainties. Although the genome graph is the main output of MinYS, it may be necessary to analyze it further to get conventional linear assemblies. This can be done either by manual inspection using interactive tools such as Bandage (20), or by enumerating all possible paths within the graph.

#### Implementation and availability

MinYS is implemented as a python3 script and available at https://github.com/cguyomar/MinYS and as a conda package in the bioconda repository. Notably, it relies on the local assembly tool *MindTheGap*, starting from version 2.1 that enables the so-called “contig mode” (https://github.com/GATB/MindTheGap). The whole pipeline presented in Figure 1 can be run in a single command line with few parameters to tune and is implemented in a modular way enabling to start or resume the analysis at intermediary steps. The most influential parameters concern the two *de novo* assembly steps (backbone and gap-filling). They are standard parameters of De Bruijn Graph based assemblers that depend mainly on the expected read depth of the targeted genome in the sample, namely the size of *k* − mers and the *k* − mer minimal abundance used to filter out sequencing errors.

#### Application to pea aphid metagenomic datasets

In this study, we applied MinYS to the assembly of the pea aphid obligatory symbiont, *Buchnera aphidicola*. We considered 50 pea aphid resequencing samples of *Illumina* 100 bp paired-end reads (21). 32 datasets are the result of sequencing a single clonal population of aphids. In addition, 18 samples sequenced from pooling 14 to 35 pea aphid individuals were also analyzed. These samples are more challenging for metagenomic assembly due to their richer symbiotic content and strain diversity (21).

#### Reference genomes with increasing levels of divergence

Reference-guided assembly was performed with 4 distinct reference genomes of *B. aphidicola* with different levels of divergence: 1) *B. aphidicola* from *A. pisum* (LSR1 accession), hereafter called *the closest*, which is the closest available assembled genome; 2) *B. aphidicola* strain G002, which was isolated from a different aphid host, *Myzus persicae*, hereafter called *distant*; 3) *B aphidicola* strain Sg, isolated from an even more distant aphid host, namely *Schizaphis graminum*, the most divergent reference analyzed and here-after called *most distant*; and 4) a synthetic genome obtained by deleting 116.4 Kb of sequences from the closest reference genome (*Buchnera aphidicola LSR1*). The later synthetic genome was generated by applying 20 deletions, whose size ranged from 300 bp to 20 Kb. The levels of divergence of these different reference genomes are supported by phylogenetic studies (22), whole genome alignments (ANI) and relative amounts of mapped reads from the real sequencing samples, compared to the clostest reference genome (see Table 1).

**Table 1.** Description of the different reference genomes for *Buchnera aphidicola* used in this study, and their levels of divergence with the targeted genome. ANI stands for Average Nucleotie Identity and is computed on the aligned portions of the genome.

|  | Name | Host | Accession | Length (bp.) | Proportion of mapped reads | ANI |
| --- | --- | --- | --- | --- | --- | --- |
| Closest | *Buchnera aphidicola* str. LSR1 | *Acyrtosiphon pisum* | NC_002528.1 | 642,011 | 100% | 100% |
| Incomplete | *Buchnera aphidicola* str. LSR1 | *Acyrtosiphon pisum* |  | 525,611 | 81% | 100% |
| Distant | *Buchnera aphidicola* str. G002 | *Myzus persicae* | NZ_CP002701.1 | 643,517 | 48% | 80.75% |
| Most distant | *Buchnera aphidicola* str. Sg | *Schizaphis graminum* | NC_004061.1 | 641,454 | 30% | 78.40% |

**Table 2.** Graph complexity metrics of the genome graphs output by MinYS for individual (top table) and pool (bottom table) samples with several reference genomes used as a guide. Values indicate the percentage of samples with the given graph characteristics.

| Individual samples | Reference genome |  |  |  | Average |
| --- | --- | --- | --- | --- | --- |
|  | Closest | Incomplete | Distant | Most distant |  |
| Assembly in a single connected component | 93.75% | 93.75% | 96.88% | 87.50% | 92.97% |
| Assembly in a single path (before comparison) | 71.88% | 71.88% | 81.25% | 81.25% | 76.57% |
| Assembly in a single path (after comparison) | 96.88% | 96.88% | 96.88% | 93.75% | 96.10% |
| Pooled samples | Reference genome |  |  |  | Average |
|  | Closest | Incomplete | Distant | Most distant |  |
| Assembly in a single connected component | 100.00% | 100.00% | 100.00% | 88.89% | 97.22% |
| Assembly in a single path (before comparison) | 11.11% | 5.56% | 5.56% | 0.00% | 5.56% |
| Assembly in a single path (after comparison) | 55.56% | 61.11% | 88.89% | 27.78% | 58.34% |

#### Dataset with simulated structural variations

To assess the ability of MinYS to recover structural variations in samples with strain diversity, we created a synthetic pea aphid sample by adding to a real sample, a subset of simulated reads from the previously described *incomplete* genome (with 20 deletions). 50X coverage of 100 bp reads were simulated with wgsim of the Samtools suite (23), using the parameters ‘-1 100 -2 100 -N 131400’.

#### MinYS parameters

MinYS was applied with the same set of parameters for all samples and reference genomes. For the assembly step, a *kmer* size of 61 was chosen, along with a *kmer* minimal abundance threshold of 10, and a minimum contig length of 400 bp. The gap-filling step was performed using version 2.2.0 of MindTheGap, with a *k* value of 51, *kmer* minimal abundance threshold of 5 and arguments *maxlength* and *max-nodes* set to 50,000 and 300.

#### Path enumeration in genome graphs

Genome graphs encompass the genomic diversity present in a sample, and therefore can often not be simply converted in a single genomic sequence. To allow the analysis of usual assembly metrics and comparisons with other assembly approaches, genome graphs were analyzed to recover a single genome sequence when only punctual polymorphism or small assembly uncertainties distinguished the different paths in the graph, or otherwise a set of genome sequences, that were representative of the different genome structures. For each connected component, all possible paths were enumerated. A clustering of those paths was performed in order to output a subset of significantly different sequences. Paths were compared using the ANI metric, as implemented in pyani (24). Two paths were considered identical if an alignment with more than 99% of sequence coverage and more than 99% of nucleotide identity could be obtained. After all the paths have been enumerated and compared, the longest resulting sequence is arbitrary chosen and considered as the representative genome sequence, to compute assembly metrics and to be compared to other approaches.

#### Comparison with other approaches

Alternative usual approaches to assemble a particular genome from metagenomic data were applied on the 50 pea aphid samples using the *distant* reference genome.

Two reference guided assembly tools were used: Mitobim (11) and Metacompass (15), which is based on Pilon(13). Mitobim was run with the ‘-quick’ parameter allowing to supply a reference genome for read baiting, and a maximal number of iterations of 31. Metacompass was used in its reference guided mode with default parameters. Alongside reference polishing using Pilon, Metacompass also uses Megahit(9) to assemble all unmapped reads. As such, the assembly returned by Metacompass is not targeted, and contains contigs representing all the sequenced genomes. To extract only the contigs associated with *B. aphidicola*, we performed a Blast alignment against the chosen reference genome, and retained the contigs with at least 50% of their length covered by Blast hits with e-values smaller than 10^−5^ were kept. The final MetaCompass assembly therefore includes the Pilon corrected sequences and the blast-filtered MegaHit contigs. Alternatively, an assembly-first approach was used for comparison. A complete *de novo* assembly was performed for each sample using Megahit (9) and *B. aphidicola* contigs were selected by a Blast alignment against the chosen reference genome, similarly to what was done for MetaCompass (at least 50% of sequence coverage and e-value smaller than 10^−5^). Similarly to what was done with MinYS, we did not include contigs smaller than 1 Kb, mainly associated with plasmid sequences.

The quality of each assembly was assessed using Quast (25) and the closest reference genome (*Buchnera aphidicola str. LSR1* from *A. pisum*). Several assembly metrics were compared, such as their length relative to the targeted genome size, the number of contigs, and the length of the largest contig. Those metrics were also computed for the backbone assembly performed at the first step of MinYS, mainly as a way to measure the relative contributions of reference-based assembly and gap-filling to the final assembly.

## Results

In this study, we applied MinYS to the assembly of the obligatory bacterial symbiont of the pea aphid holobiont, *B. aphidicola* (640 Kb). We considered a large set of pea aphid resequencing samples of *Illumina* 100 bp paired-end reads, that have already been studied in a previous work, in which the microbiota of each aphid sample was detailed (21). Among the 50 datasets, 32 are the result of sequencing one single aphid clone, and 18 are each the result of sequencing a pool of individuals from a same population. The number of reads per dataset is on average 84 (resp. 198) millions for individual (resp. pool) sequencings, with an average coverage of 628X (resp. 3,694x) for the symbiont *B. aphidicola* genome. Notably, in such datasets, more than 90% of the reads originate from the insect host, and are not relevant when focusing on symbiont genomes. This particular fact motivates the choice of a targeted assembly approach, which does not require to assemble pea aphid reads.

### Single contig assembly of the targeted genome using increasingly distant reference genomes

MinYS was first applied to the set of 32 resequenced aphid clones using several reference genomes. Assembly statistics are shown in Table 3. Overall, the *Buchnera aphidicola* genome was very well assembled in most cases: for all samples but one the targeted genome is assembled in one single contig whose size is comparable to the expected genome size. Overall, the sequence accuracy witnessed for all the assessed methods was satisfying and in range with the expected divergence to the closest reference genome.

**Table 3.** Assembly metrics of different assembly approaches using the *distant* reference genome. For each metric, the median value obtained over all individual or pool samples is given. Extended statistics are available in Supplementary Table 1.

| Tool | Length / reference length (%) |  | Nb. contigs |  | Largest contig (kb) |  | Runtime (h) |  |
| --- | --- | --- | --- | --- | --- | --- | --- | --- |
|  | Individuals | Pools | Individuals | Pools | Individuals | Pools | Individuals | Pools |
| MinYS | 100 | 101.0 | 1.0 | 1 | 642 | 646 | 0.64 | 2.34 |
| Metacompass | 108 | 113 | 54.5 | 90 | 642 | 310 | 6.69 | 29.16 |
| Megahit | 100 | 102 | 1.5 | 11.5 | 642 | 366 | 5.00 | 15.98 |

As MinYS relies on a reference genome to perform the assembly, we compared the assembly results using increasingly distant reference genomes (Table 1). Figure 2 presents the results obtained from the different considered genomes, and shows the contributions of the first (reference based mapping and assembly) and second (gap-filling) steps of MinYS. The main result is that using an incomplete or distant genome did not affect the assembly quality: both assembly length and contiguity are similar to those obtained using the closest genome. Using the incomplete genome, including 20 deletions between 300 bp and 20 Kb, did not hinder the assembly with all missing regions from the reference genome being assembled, illustrating how MinYS is able to recover large previously unknown regions of the genome. Only the use of the most distant genome (*B. aphidicola* from *S. graminum*) decreased the completeness and contiguity of some assemblies. In that extreme case, still 88% of samples were assembled in a single contig, and 90% of the assemblies had their length greater than 98% of the targeted genome length.

**Fig. 2.**
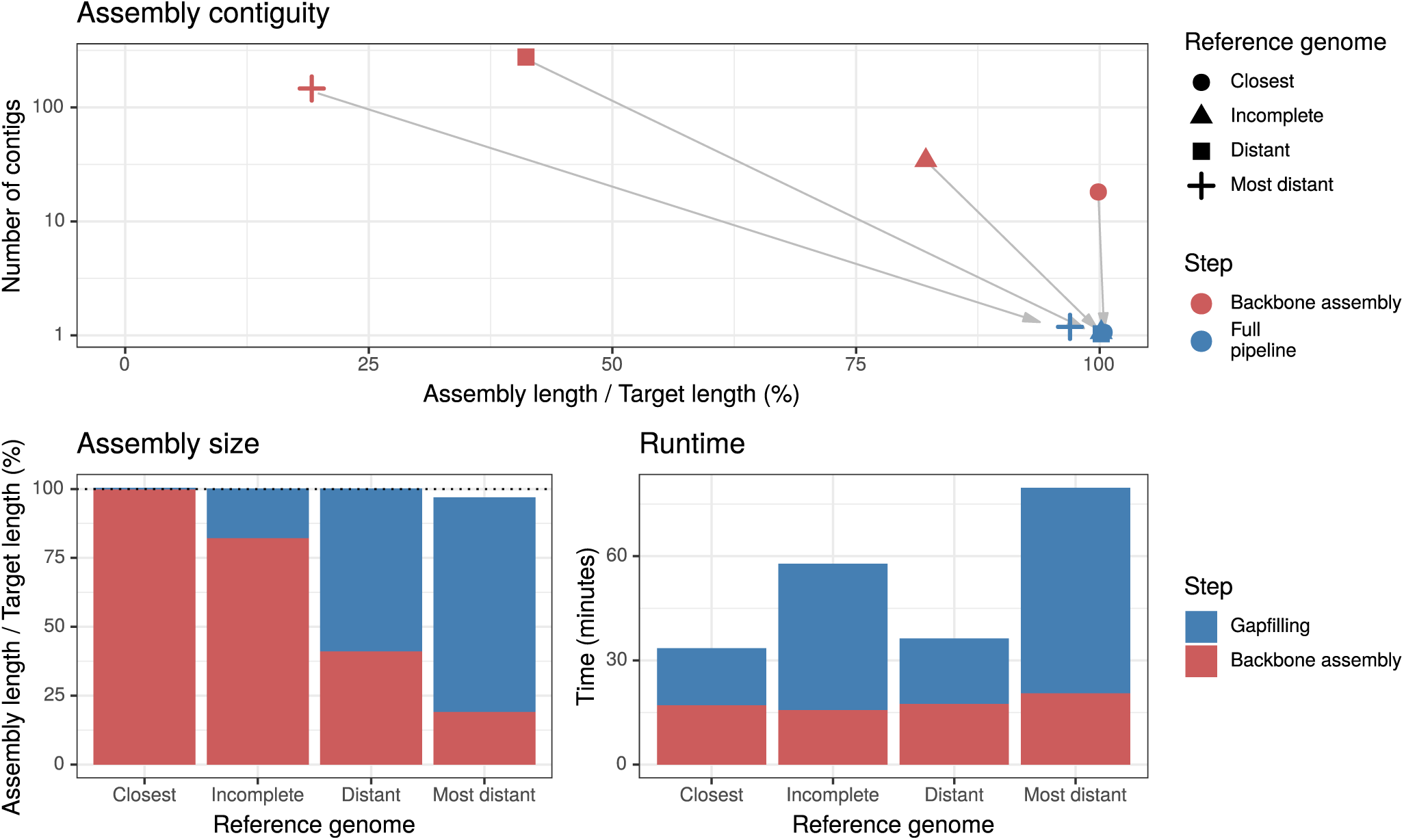
Effect of the level of divergence of the reference genome on MinYS assembly statistics (assembly contiguity on the top plot, assembly completeness on the bottom left plot, and running time on the bottom right plot) and on the relative contribution of the first backbone assembly step (in red) and of the gap-filling step (blue). The plots show the average values for the 32 individual samples.

However, using more distant reference genomes affected the MinYS pipeline and therefore its intermediate results. Using more distant reference genomes resulted in fewer reads mapped and assembled during the first step of the pipeline, and therefore a more partial initial backbone assembly. The fraction of the genome assembled during the first mapping and assembly step is 99% for the *closest*, 47 % for the *distant* and 24 % for the *most distant* reference genome. As a consequence, the gap-filling step is essential when using more distant genomes. Since final assembly results are comparable, this shows that this step successfully recovered the genome portions highly divergent or missing from the reference genome. Accordingly, the running time is increased when a remote reference is used, mainly due to the increase of time devoted to gap-filling.

### Assembly of pooled samples with a higher strain diversity

MinYS was then applied to an additional set of 18 sequencing samples of pooled aphids (with 15 to 34 aphid individuals per pool). An important feature of MinYS is to output a genome graph, that can represent several putative assemblies. In the case of pooled samples, the genome graph output by MinYS encompasses more diversity than in individual samples, as shown by the higher number of enumerable paths in these graphs (see Table 2). While 77% of the individual samples were assembled in a single path, it was the case for only 6% of the pooled samples. This shows that MinYS is able to recover the logically higher level of strain diversity present in pooled samples. Interestingly, most alternative paths in these pooled samples differed only by punctual polymorphism and did not encompass large structural variations. Clustering the enumerated paths for each sample with a 99 % sequence identity threshold resulted in one single representative sequence for 58 % of the pooled samples. Noteworthy, using the most distant reference genome as a primer resulted in more complex graphs and pushed to its limits this post-processing step of path clustering. In that case, four samples were not considered due to excessively high computational time for the comparison of sequence paths. When more than one representative sequence was generated, the longest path was arbitrary selected, explaining why the assembled length is slightly higher in pooled samples (see Table 3).

### Assembly of coexisting structural variants

MinYS was applied to a pea aphid sample in which simulated reads from a rearranged *B. aphidicola* genome were added to a real pea aphid re-sequencing sample, simulating the coexistence in a metagenomic dataset of two strains with structural variations (here 20 deletions with size between 300 bp and 20 Kb). In the resulting genome graph, shown in Figure 3, 17 out of the 20 simulated deletions were fully recovered, with both the deleted and complete versions of the genome assembled. Extracting the longest path from the graph resulted in a single contig of 641.5 Kb, compared to the 642 Kp of the closest reference genome. Similarly, the shortest path extracted from the graph was 526.4 Kb long, compared to 525.6 Kb for the simulated deleted genome. The longest structural variations (up to 20 kb) were all successfully recovered. Only two small variations, of 300 and 500 bp, were missing from the graph.

**Fig. 3.**
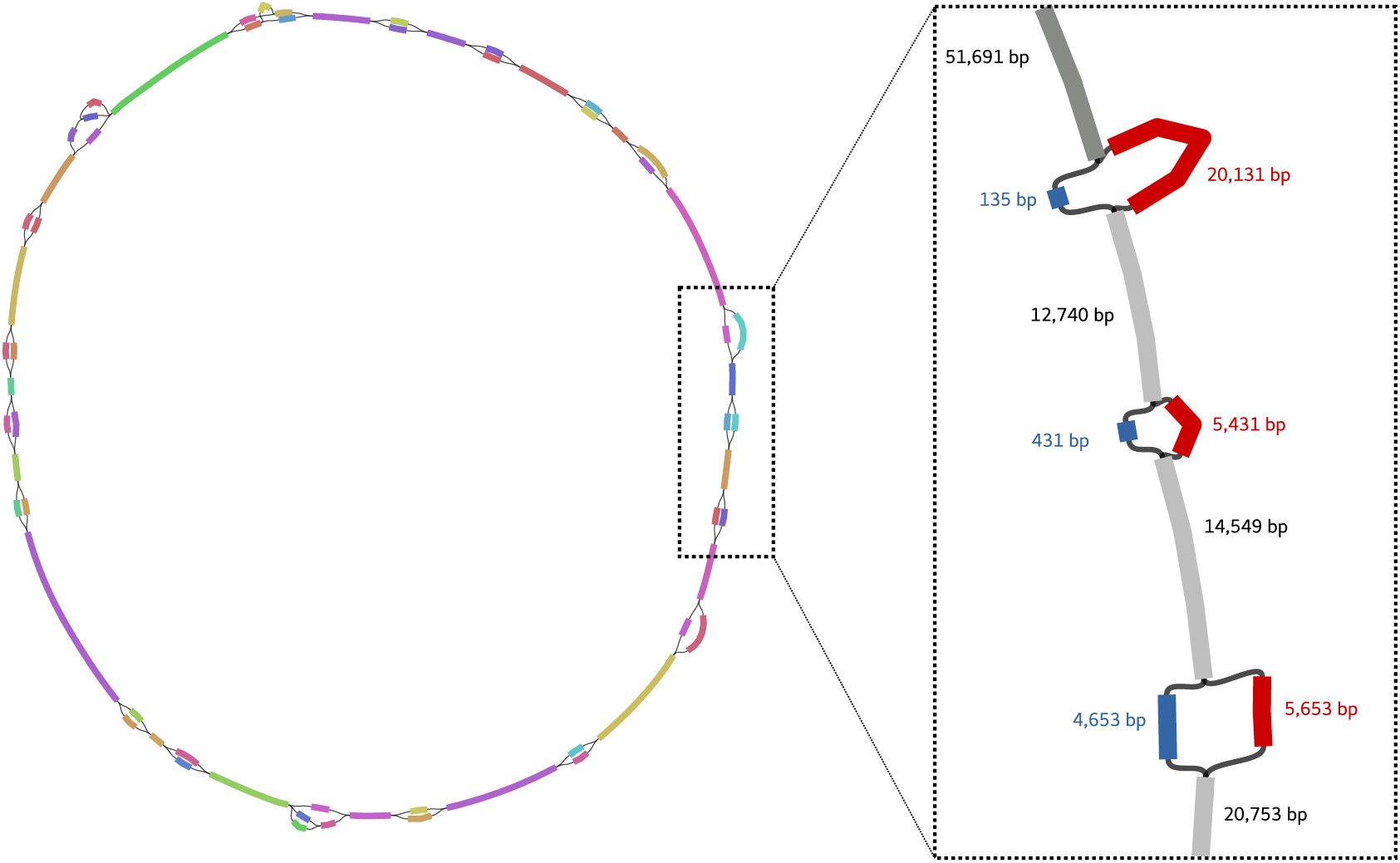
View of the genome graph generated by MinYS with a sample containing two coexisting strains of *Buchnera aphidicola* differing by 20 structural variants. The graphical representation was obtained with the assembly graph visualisation tool, Bandage, where each node is represented by a coloured rectangle whose size is related to the node sequence size.

### Comparison with other approaches for targeted genome assembly

MinYS assemblies were compared to the results of other reference based approaches (MitoBIM (11), Metacompass (15)) and a *de novo* metagenomic assembly (Megahit (9)) followed by blast-based contig filtering. Results using the *distant* reference genome are shown in Table 3.

MitoBIM was designed for reference guided assembly of mitochondrial genomes by read baiting and mapping (11). When applied to a pea aphid metagenomic sample, it reached its maximal number of iterations (31 iterations) in 6 hours, and was unable to assemble a complete genome. As such, it does not scale well with genomes larger than mitochondrial genomes, and was not considered as an alternative to MinYS in that context.

Metacompass first polishes the supplied reference genome sequence with mapped reads using Pilon (13), and then assembles in a *de novo* manner the remaining unmapped reads using Megahit. To recover, among the Megahit contigs, only the ones originating from the genome of interest, an additional step of mapping to the reference genome is therefore necessary and was performed here with blast. The first step was insufficient to return satisfying assemblies with that level of divergence. Considering only the polished sequences returned by Pilon resulted in far from complete assemblies, with an average length of 6% of the targeted length for individual samples. Therefore, the assemblies described in Table 3 are dominated by *de novo* assembled contigs, mitigating the referenced-based approach of Metacompass. Overall, obtained assemblies were more fragmented and longer than expected, due to potential redundancy between polished and *de novo* contigs.

Finally, we compared MinYS to a full metagenomic assembly approach with Megahit, followed by blast-based contig filtering. On individual samples, MegaHit performs on par with MinYS, resulting in complete one-contig assemblies in most cases. However, when applied to pooled samples with more polymorphism, Megahit does not perform as well, with more and shorter contigs (the median size for the largest contig is 57% of the expected genome length, compared to 101% with MinYS). An explanation for this could be that highly polymorphic regions may be assembled by Megahit into distinct contigs that break the contiguity, while MinYS often represents them as bubbles in the genome graph.

Notably, the metagenomic assembly approach with Megahit performed badly compared to MinYS in the experiment with co-existing structural variations in the metagenomic dataset. It resulted in a 38 contigs assembly, with a length of 646 Kb and a N50 of 44.5 Kb, whereas several structurally different strains all assembled in a single contig can be extracted from the genome graph output by MinYS. This highlights the difficulty of *de novo* assembly to deal with structural diversity in metagenomic samples.

Importantly, MinYS is also significantly faster than both Megahit and Metacompass. The average runtime of MinYS full pipeline is 38 minutes for individual samples and 6.5 hours for pooled samples, which is respectively 10 times and 12 times inferior to Megahit runtimes. Indeed, Megahit produces contigs not only for the targeted organism, but in this case for the insect host *A. pisum* and all its secondary symbionts. This highlights the importance of a tool such as MinYS, to efficiently recover specific genomes of interest from metagenomic data.

## Discussion and conclusion

### *De novo* and reference guided assembly: the best of both worlds ?

When working on microbial communities, focusing on the assembly and study of a particular organism can be especially relevant. For biologists, understanding these communities can be achieved by focusing on some key players with particular functions or ecological impacts. For bioinformaticians, assembling a single genome can be significantly less challenging than a whole community, especially in the context of symbiotic associations where reads originating from the organism of interest are a minority within host-dominated sequence data. Yet, existing tools for targeted assembly are usually not suited for metagenomics data, or rely on reference sequence correction, which is unable to capture novel sequences absent from the reference genome.

MinYS is a novel method leveraging the benefits of both reference based and *de novo* assembly. By using a reference genome as a primer for the assembly, MinYS significantly reduces the computational burden of genome assembly from a metagenomic sample. In the first step, the assembly is restricted to a subset of the reads, and is therefore usually straightforward. During gap-filling, although the whole readset is used, performing local assembly between specific seed and target kmers is also less demanding than complete metagenomic assembly. In addition, as long as trustworthy contigs are assembled in the first place, this local gap-filling is less prone to include contaminant sequences. This makes the assembly both more reliable and faster. In this context of mining a particular symbiont genome out of a host-dominated sequencing dataset, MinYS was up to 12 times faster than a full metagenomic assembly.

MinYS obtained also better contiguity statistics, with the genome of *Buchnera aphidicola* being assembled in one circular sequence in most cases. Admittedly, this organism has a small and simple genome containing few repeated sequences, enabling to obtain full length assemblies with short read data. This may not be the case for many other organisms, but this kind of genome is a typical use case of our approach which was designed for symbiotic community studies. Moreover, although apparently simple, our analyses have shown that depending on the sequencing context, the assembly task may not be so straightforward, as highlighted by the lower contiguity of Megahit assemblies when strain diversity was present in the sample. In this context, MinYS proved significant improvements in assembly contiguity with respect to other approaches. Although full metagenomic assembly also delivers contigs for the other organisms of the community, we consider that the speed increase along with the contiguity improvement make of MinYS a worthy alternative to analyze particular components of microbial communities.

### MinYS performs reference based assembly, but not reference dependant assembly

Alternative reference-based approaches usually rely on read mapping followed by a correction of the existing reference genome. On the contrary, every sequence returned by MinYS has been assembled from the metagenomic reads. This reduces the bias due to the choice of a reference genome. Thanks to the gap-filling step, it is also possible to assemble *de novo* regions too distant or absent in the reference genome to be captured by reference-based approaches. These regions represented up to 80% of the targeted genome in the most extreme case considered here.

Remarkably, MinYS proved to be particularly robust with respect to the level of divergence with the chosen reference genome. Thanks to its two step approach, MinYS can adapt to different divergence levels. When using a close reference genome, the first mapping based phase is predominant, and the assembly is fast. On the opposite, when using more distant reference genomes, the gap-filling step is essential to recover regions missing or too divergent compared to the reference. MinYS can deal with both a high level of divergence and the presence of large novel inserted sequences. When based on an incomplete reference genome, it was able to recover all the missing sequences (including up to 20 Kb consecutive missing parts).

In this work, a single set of parameters was chosen for all samples, regardless of divergence or sample coverage. However, MinYS is flexible and also allows to use more or less conservative parameters to adapt to particular use-cases, for instance to accommodate with higher levels of divergence. Overall, although MinYS relies on a reference genome as a primer for the assembly, its hybrid approach allows to use distant reference genomes, while still being able to return full length assemblies.

### MinYS assembles and represents genomic diversity

Species in metagenomic samples may not be homogeneous populations sharing the same genome. As highlighted by the results of Megahit on pooled samples, a high level of polymorphism within a sample affects the contiguity of the assembly. Genomic *loci* showing significant nucleotide diversity are assembled in separate contigs instead of single consensus contigs and they fragment the assembly. In MinYS, such *loci* are deliberately represented as distinct nodes in the genome graph, but are still connected together. As such, the genome graph encompasses different assemblies, and its complexity is related to the genomic diversity in the sample. Accordingly, pool sequencing samples, supposed to shelter a greater microbial diversity, were assembled into more complex graphs. These graphs contain biological meaningful information that surpasses the “one species- one genome” paradigm which has been lately increasingly criticized (26, 27). As a consequence, many novel tools are being developed to use such genome graphs as a reference rather than a conventional linear sequence, for instance for read mapping or variant genotyping (28, 29).

Genome graph representation is also a powerful way to represent structural variations in metagenomic samples. In the context of two coexisting strains differing by large structural variations, MinYS was able to reconstruct both strains as alternative paths in the genome graph, while Megahit yielded a fragmented assembly.

Genome graphs are an opportunity to better represent genomic diversity in metagenomic samples, but are also challenging objects to study. The main output of MinYS is a genome graph, that may require some post-processing to output one or several representative sequences. Here, all possible paths in the graphs were enumerated and compared to each other to choose a single representative genome. Additionally to the computational burden of such an enumeration, many of these paths may not represent real present strains. Therefore, an interesting but far from trivial perspective of this work would be to map back the paired-reads on the graph in an attempt to phase the different alleles, and then offer a more accurate representation of present genome strains in the sample.

## Acknowledgements

Computations have been made possible thanks to the resources of the Genouest infrastructure. The authors warmly thank Guillaume Rizk for interesting discussions and valuable help on implementation with the GATB library.

