## Supplementary material for "MinYS: Mine Your Symbiont by targeted genome assembly in symbiotic communities"

#### 1 Contig mode of MindTheGap

The gap-filling step of MinYS is based on a module of the software *MindTheGap*, originally developed for the detection and assembly of long insertion variants [**?**]. The original *fill* module of *MindTheGap* takes as input a read set and a set of insertion variant breakpoints, of the form of two kmers, the left and right kmers adjacent to each insertion site. It first builds a de Bruijn graph of the entire input read set, and then performs a local assembly between the left and the right kmers of each insertion site, by looking for all the paths in the De Bruijn graph starting from the left (source) kmer and ending in the right (target) kmer.

In this work, we took advantage of this module of *MindTheGap* and adapted it to the problem of simultaneous gapfilling between multiple contigs. It has been modified to make possible the gapfilling between a source kmer and **multiple** target kmers, enabling the "all versus all" gapfilling within a set of contigs with only a linear increase of the runtime (compared to a quadratic increase for a naive "all versus all" gapfilling as if using *MindTheGap* in its original breakpoint mode).

The resulting algorithm is presented in Figure 1. A seed kmer is extracted at the end of each contig and its reverse-complement, resulting in a set of $2n$ source kmers for $n$ contigs. Similarly, a set of $2n$ target kmers is extracted at the beginning of each contig and its reverse-complement. For each seed kmer, a contig graph is created by starting from the seed kmer and performing a breadth first traversal of the *De Bruijn* graph representation of the whole readset. Contigs are consensus sequences returned by removing graph motifs such as bubbles (SNPs) and tip-ends (errors). In the contig graph, contigs are nodes, and edges represent the existence of a $k-1$-mer overlap between two contigs. The algorithm creation of the contig graph is similar to the one used in *Minia* [**?**]. The traversal is stopped when the graph becomes too large (total assembled nucleotides) or too complex (number of contigs), following user-defined parameters. Importantly, if one of the target kmers is found during the contig graph construction, that contig is not extended further, avoiding redundant contig assembly, and saving time and memory. After the contig graph has been built, target kmers are searched within this contig graph, and gapfilling sequences are built, by retraversing the contig graph from each target kmer to the source kmer. For each seed-target couple, if several sequence solutions are returned, redundant solutions above a 95% identity threshold are removed. Thanks to this multi-target version of the algorithm, only $2n$ contig graph constructions are necessary to search all possible gapfilling sequences between all pairs of contigs, instead of $n^2$ with the naive approach.

The whole process is parallelized by dispatching the $2n$ starting kmers to different threads. The main output is a genome graph in the GFA format (Graphical Fragment Assembly, `https://github.com/GFA-spec/GFA-spec`), containing all input contigs and their gap-filling sequences as nodes, together with their overlap relationships as edges.

All these changes have been implemented in the so-called *contig mode* of *MindTheGap* (since version 2.1), whose input is a read set together with a set of contigs and output is a sequence graph in GFA format.

Figure 1: **Gapfilling a set of contigs using *MindTheGap* fill module**.
a) Seed and target kmers are extracted from the 3 input contigs, resulting in 2 sets of 6 kmers, seed (red) and target (blue) ones. b) A graph of contigs is built starting from the right seed kmer of contig A. Extension is stopped when a target kmer of another contig if encountered, or a maximum assembly size is reached. c) This results in 3 gapfilling sequences starting from contig A right seed, 2 gapfilling sequences joining contig B, and one contig C.

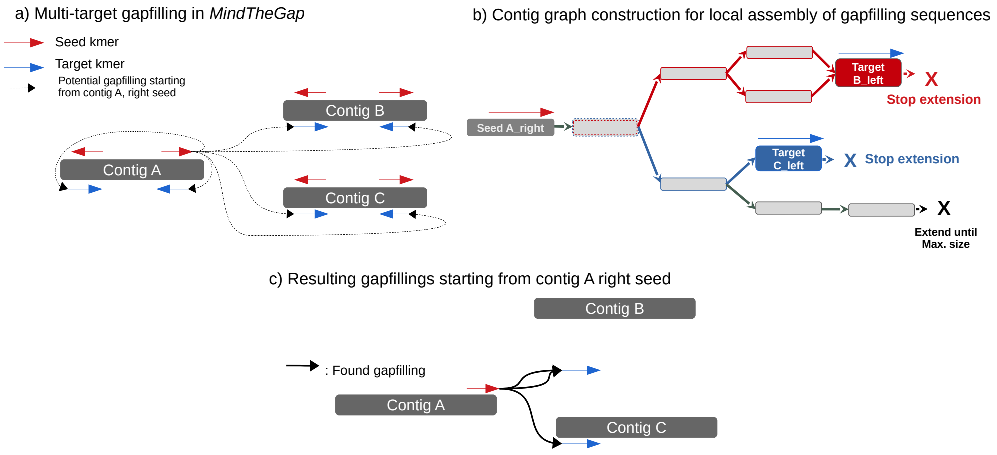


#### 2 Graph simplification

Figure 2: **Graph simplification applied to gap-filling output**.
a) Reciprocal gapfillings with more than 95% identity are merged. b) Shared sequences between gapfillings originating from or leading to a same contig are merged to reduce sequence redundancy c) Simple linear paths, with no branching nodes, are merged into a single node.

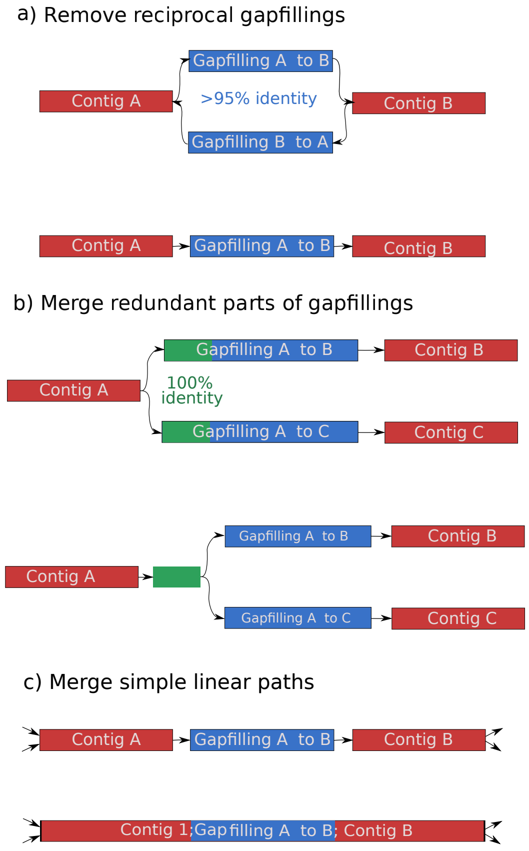

### 3 Full assembly statistics

Table 1: Assembly results in 50 pooled and individual samples for MinYS and Megahit. Values are : median (minimum-maximum)

| | Length / reference length (%) | | Nb. contigs | | Largest contig (kb) | | Runtime (h) | |
|---|---|---|---|---|---|---|---|---|
| Tool | Individuals | Pools | Individuals | Pools | Individuals | Pools | Individuals | Pools |
| **Closest** | | | | | | | | |
| MinYS | 100 (100-113) | 101 (100-109) | 1 (1-2) | 1 (1-1) | 642k (437k-727k) | 648k (642k-703k) | 0.55 (0.18-1.2) | 2.29 (1.33-5.42) |
| Megahit | 100 (100-102) | 102 (101-104) | 1 (1-9) | 15 (5-30) | 642k (348k-643k) | 366k (86k-642k) | 5 (2.14-10.76) | 15.98 (2.97-34.44) |
| **Incomplete** | | | | | | | | |
| MinYS | 100 (100-101) | 101 (100-107) | 1 (1-2) | 1 (1-1) | 642k (437k-646k) | 646k (642k-686k) | 0.93 (0.22-1.83) | 3.32 (1.26-8.23) |
| Megahit | 100 (100-100) | 100 (90-101) | 1 (1-3) | 4.5 (1-16) | 642k (348k-643k) | 366k (86k-642k) | 5 (2.14-10.76) | 15.98 (2.97-34.44) |
| **Distant** | | | | | | | | |
| MinYS | 100 (95-114) | 101 (100-109) | 1 (1-2) | 1 (1-1) | 642k (437k-730k) | 646k (642k-698k) | 0.64 (0.19-0.99) | 2.34 (1.27-4.51) |
| Megahit | 100 (100-102) | 102 (101-104) | 1.5 (1-7) | 11.5 (5-30) | 642k (348k-643k) | 366k (86k-642k) | 5 (2.14-10.76) | 15.98 (2.97-34.44) |
| **Most distant** | | | | | | | | |
| MinYS | 100 (57-114) | 101 (1-119) | 1 (1-3) | 1 (1-1) | 642k (262k-729k) | 648k (5419-765k) | 0.87 (0.18-3.54) | 2.63 (1.57-8.18) |
| Megahit | 100 (100-102) | 102 (101-104) | 1.5 (1-8) | 13 (5-31) | 642k (348k-643k) | 366k (86k-642k) | 5 (2.14-10.76) | 15.98 (2.97-34.44) |
